# First report of reference guided genome assembly of Black Bengal goat (*Capra hircus*)

**DOI:** 10.1101/603266

**Authors:** Amam Z. Siddiki, A. Baten, M. Billah, MAU. Alam, KSM. Shawrob, S. Saha, M. Chowdhury, A.H. Rahman, M. Stear, G. Miah, M. Kumkum, M.A. Hossain, AKM. Mollah, M.S. Islam, MKI. Khan

**Affiliations:** Genomics Research Group, Faculty of Veterinary Medicine, Chittagong Veterinary and Animal Sciences University (CVASU), Chittagong, 4225, Bangladesh; AgResearch, Palmerston North 4410, Private Bag 11008, New Zealand; Department of Computer Science and Engineering, Bangladesh University of Engineering and Technology (BUET), Dhaka 1000, Bangladesh; AgriBio, Department of Animal, Plant and Soil Sciences, School of Life Sciences, La Trobe University, Bundoora, Victoria 3083, Australia; Department of Genetics and Animal Breeding, Faculty of Veterinary Medicine, Chittagong Veterinary and Animal Sciences University (CVASU), Chittagong, Bangladesh; Department of Biological Sciences, Asian University for Women (AUW), Chittagong, Bangladesh

**Keywords:** Black Bengal goat, *Capra hircus*, Whole Genome sequence, Assembly, Annotations

## Abstract

**Objectives:** Black Bengal goat (*Capra hircus*), a member of the Bovidae family with the unique traits of high prolificacy, skin quality and low demand for food is the most socioeconomically significant goat breed in Bangladesh. Furthermore, the aptitude of adaptation and disease resistance capacity of it is highly notable which makes its whole genome information an area of research interest.

**Data description:** The genomic DNA of local (Chittagong, Bangladesh) healthy Black Bengal goat (*Capra hircus*) was extracted and then sequenced. The *de novo* assembly and structural annotations are being presented here. Sequencing was done using Illumina sequencing platform and the draft genome assembled is about 3.04 Gb. 26458 Genes were annotated using Maker gene annotations tool which predicted BUSCO Gene models. Universal Single Copy Orthologs refer 82.5% completeness of the assembled genome.

## Objective

Black Bengal goat (BBG) belongs to the Bovidae family and found throughout Bangladesh, West Bengal, Bihar, and Orissa regions of northeastern India. It is estimated that more than 90% of the goat population in Bangladesh comprised the Black Bengal, the remainder being Jamunapari and their crosses [1]. Higher prolificacy, fertility, resistance against common diseases, adaptability to the adverse environmental condition, early maturity, seasonality and superiority in the litter size are some of the outstanding features of BBG. Besides, it produces excellent quality flavored, tender and delicious meat with low intramuscular fat and fine skin of extraordinary quality for which there is tremendous demand all over the world [1, 2]. Moreover, it plays a vital role in the economy of Bangladesh by contributing 1.66% of the GDP (Gross Domestic Product) (DLS, 2017).

Fortunately, the market demand of Black Bengal goat is emerging. This gives breeders of original/rare breeds an opportunity to expand the stock and preserve its genetic diversity. One of the primary goals in managing goat populations is to maintain high-level genetic diversity and low-level inbreeding. To estimate the future breeding potential of a goat breed, it is necessary to characterize the genetic structure and evaluate the level of genetic diversity within the breed. Moreover, a long term genetic approach can be used to improve the spectacular economic characteristics of BBG [3].

Therefore, the genetic characterization of the entire BBG genome is essential in characterizing its economic traits as well as adaptive capability. With the availability of whole genome sequence, the targeted areas for genetic improvements are now: goat prolificacy, growth rate, meat quality, skin quality, disease resistance, and survivability. A complete and accurate reference to the goat genome is an essential component of advanced genomic selection of product characteristics.

## Data description

At first, healthy Black Bengal Goats (BBG) without known genetic diseases were selected for blood collection. Genomic DNA from each animal was isolated from the EDTA-blood, using the Addprep genomic DNA extraction kit (South Korea) (detailed methodology in Data File 1, Table 1). The quality and quantity of the DNA were assessed by the Qubit fluorometer (Invitrogen, Carlsbad, CA, USA) and Infinite F200 microplate reader (TECAN), according to the manufacturer’s instruction. The status of the DNA was visually inspected by 0.8% agarose gel electrophoresis. Purified genomic DNA was sent for library preparation (detailed methodology in Data File 1, Table1) and WGS sequencing at Beijing Genomic Institute (Beijing, China). A total of 40 Gb (14-fold) of subread bases with a read length of 150 bp were generated using next-generation sequencing (NGS) technology on an Illumina HiSeq 2500 platform (detailed methodology in Data File 1, Table 1).

**Table 1:**
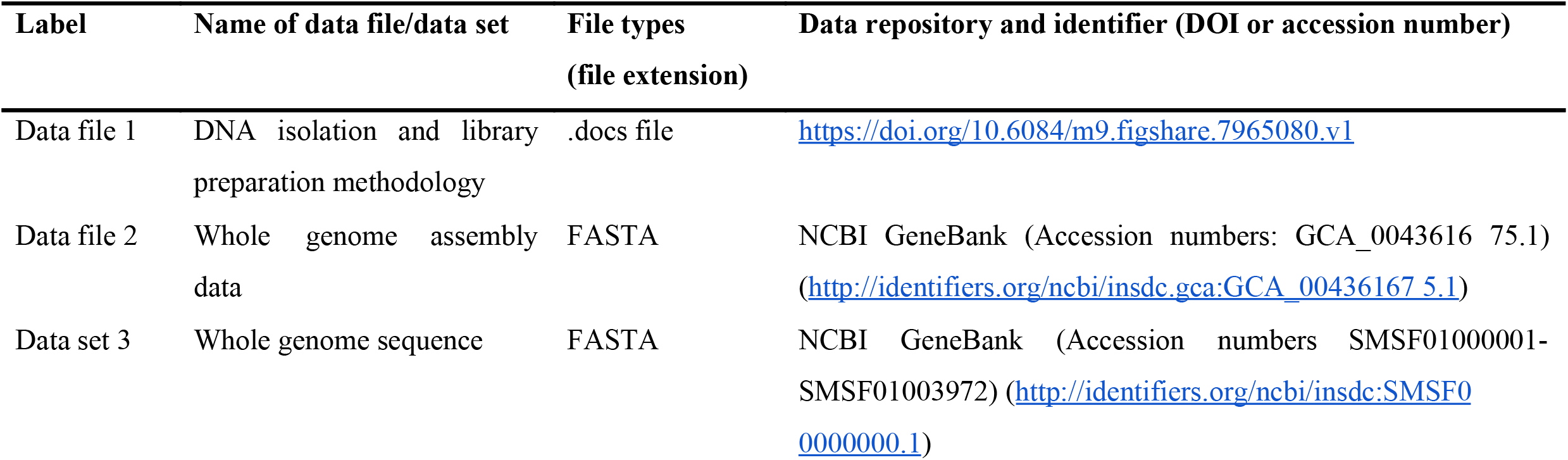
Overview of data files/data sets.

The quality of the reads was checked by using FastQC [4] and compared with the reference genome (San Clemente breed) ARS1 (GCA_001704415.1) which significantly improved the quality and continuity of the genome. As a result, the N50 increased from 87,277,232 Kb (kilobase pair) to 102,339,471 Kb and the total number of scaffold decreased from 29,907 to 3,972. Subsequently, for de novo assembly we used ABySS v. 2.1.5 assembler [5]. Furthermore, by using ABACAS v.1.3.1 and comparing with the reference genome; arranging, ordering, and orientation of the assembled genome was done [6]. The genome assembly data has been deposited in the NCBI GenBank under the Accession numbers GCA_001704415.1(Data file 2; Table 1). The final assembled genome size of *Capra hircus* is now 3.04Gb (Gigabase pair) 82.5% completeness was revealed by BUSCO [7] analysis. Moreover, a considerably better N50 size and lower scaffolds number suggested a higher quality of the genome. The whole genome sequence data has been submitted in the NCBI GenBank under the Accession number SMSF01000001-SMSF01003972 (Data file 3; Table 1).

Consequently, to do the structural annotations Maker ver 3.0 pipeline was applied [8]. In the genome, 41.77% of GC content was observed. The latest version of the repbase database identified 31.85% repeat elements by using Repeatmasker [9]. After applying MAKER gene annotation pipeline 26458 genes were predicted which was based on both Denovo and reference guided. Finally, InterProScan [10] predicted 12589 genes out of 26458 genes, and 8173 genes have at least one GO term.

## Limitations

The number of unassembled regions in the genome is 3943.

## Abbreviations

BBG: Black Bengal Goat;
GDP: Gross Domestic Production;
EDTA: Ethylene diamine tetra-acetic acid;
DNA: Deoxyribonucleic acid;
WGS: Whole Genome Sequencing;
BUSCO: Benchmarking Universal Single-Copy Orthologs;
ABACAS: Algorithm-based automatic contiguation of assembled sequences;
Gb: Gigabase pair;
Mb: Megabase pair;
Kb: Kilobase;
bp: Base pair;
GO: Gene ontology;;
SNAP: Semi-HMM-based Nucleic Acid Parser.

## Declarations

### Ethics approval and consent to participate

The experiments discussed in this investigation were approved by the Institute Review Committee of Chattogram Veterinary and Animal Sciences University

### Consent for publication

Not applicable

### Availability of data material

The genome sequence information has been accessible at DDBJ/ENA/GenBank under the Accession numbers SMSF01000001-SMSF01003972 and the assembled genome at GCA_001704415.1. The version reported in this paper is the first version, SMSF00000000.1

### Authors’ contributions

All authors contributed equally

## Acknowledgments

Authors concede the support of Beijing Genomic Institute for the sequencing service and that of Southern Cross University, Lismore, Australia for the computational support.

